# Dilp8 requires the neuronal relaxin receptor Lgr3 to couple growth to developmental timing

**DOI:** 10.1101/017053

**Authors:** Andres Garelli, Fabiana Heredia, Andreia P. Casimiro, Andre Macedo, Catarina Nunes, Marcia Garcez, Angela R. Mantas Dias, Yanel A. Volonte, Takashi Koyama, Alisson M. Gontijo

**Author notes:** Correspondence to (A.G.); (A.M.G.). These authors contributed equally to this work (Andres Garelli, Fabiana Heredia, Andreia P. Casimiro).

## Abstract

How different organs in the body sense growth perturbations in distant tissues to coordinate their size during development is poorly understood. Here, we mutated an invertebrate orphan relaxin receptor, the *Drosophila Lgr3*, and found body asymmetries similar to those found in insulin/relaxin-like peptide 8 (*dilp8*) mutants, which fail to coordinate growth with developmental timing. Indeed, mutation or RNAi against *Lgr3* suppresses the delay in pupariation induced by imaginal disc growth perturbation or ectopic Dilp8 expression. By fluorescently-tagging the endogenous Lgr3 protein and performing CNS-specific RNAi, we find that Lgr3 is expressed and required in a novel subset of CNS neurons to transmit the peripheral tissue stress signal, Dilp8, to the neuroendocrine centers controlling developmental timing. Our work sheds new light on the function and evolution of relaxin receptors and reveals a novel neuroendocrine circuit responsive to growth aberrations.

## Main text

How different organs in the body sense growth perturbations in distant tissues to coordinate their size and differentiation status during development is poorly understood^1,2^. We have previously discovered a hormone in *Drosophila*, the insulin/relaxin-like peptide Dilp8, which ensures organ and body size coordination^3^. In developing larvae, Dilp8 is produced and secreted from abnormally-growing imaginal discs. Its activity transiently delays the onset of metamorphosis by inhibiting the biosynthesis of the major insect molting hormone ecdysone by the prothoracic gland, a part of a compound endocrine structure called the ring gland^3,4^ (Fig. 1A). Loss of *dilp8* uncouples the endocrine communication between imaginal discs and the prothoracic gland, making *dilp8* mutants susceptible to uncoordinated disc growth. This results in an increase in random deviations from bilateral symmetry [fluctuating asymmetry (FA)], measurable at the population level^3^. These findings have placed Dilp8 as a central player in the interorgan communication system that mediates plasticity to promote developmental stability in *Drosophila*. However, which molecule(s) and tissue(s) sense and/or transmit this abnormal growth signal to the prothoracic gland are unknown.

**Figure 1:**
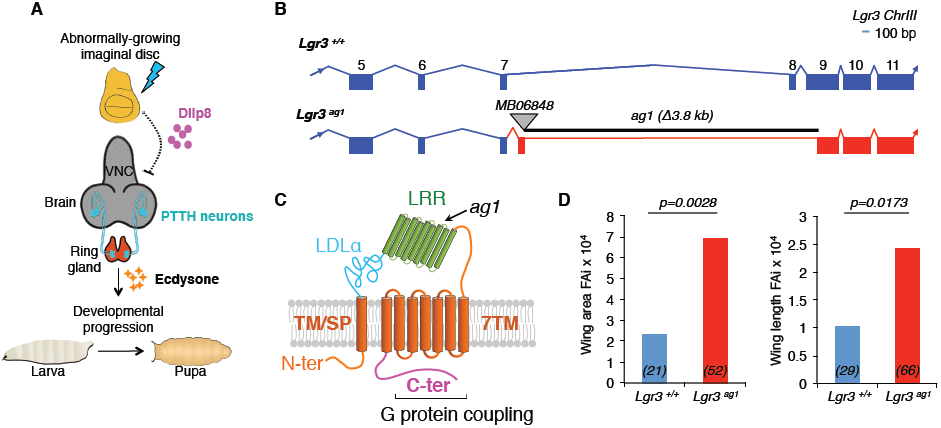
**Mutation in the *Drosophila* relaxin receptor Lgr3 leads to increased fluctuating asymmetry. A,** A neuroendocrine pathway coupling growth and developmental timing^3,4^. Scheme adapted from Halme *et al.*^9^. B, Remobilization of Minos element *MB06848* positioned between exons 7 and 8 of the *Lgr3* locus on Chromosome III generated a 3.8-kb deletion, named *Lgr3^ag1^*. **C,** Scheme of the predicted protein structure of the wild-type Lgr3 protein and the truncated Lgr3^*ag1*^ protein based on vertebrate relaxin receptor structure data*^5^*. Major domains are depicted: Low-density lipoprotein receptor domain class A (LDLa), Leucine-rich repeat (LRR), and seven transmembrane (7TM). The first TM/signal peptide (SP) domain is not predicted to be cleaved. The approximate region where *ag1* mutation truncates the Lgr3 protein is depicted. **D,** Bar graphs of the FAi*^3^* of the area (left panel) and length (right panel) of the wing pairs of the genotypes indicated. Numbers (*N*) of the wing pairs scored.

Type C1 Leucine-rich repeat-containing G-protein coupled receptors (Lgrs) are a highly conserved protein family in metazoans that act as receptors for insulin-like peptides of the relaxin subfamily in vertebrates, where they play diverse roles in tissue homeostasis and remodeling, behavior and reproduction^5^. In invertebrates, however, they are considered orphan receptors and their biological function is a mystery because invertebrates are thought to lack bona fide relaxin peptide homologues^5^-^7^. The *Drosophila* genome encodes two orphan receptors, Lgr3 and Lgr4, with clear homologies to vertebrate relaxin receptors [∼45% and ∼40% sequence similarity to human, RXFP1/2, respectively^6,7^]. To start untangling the function of these relaxin receptors, we remobilized an MB Minos element^8^ to generate a mutant for the *Drosophila Lgr3* (Fig. 1B). The obtained imprecise excision allele was named *Lgr3^ag1^.* Three precise excisions (*Lgr3^+/+^*) were also retained and used as genetic background controls. The *Lgr3^ag1^* imprecise excision (Supplementary Fig. 1A-D) consists of a 3.8-kb deletion that completely removes exon 8 and partially removes exon 9 (Fig. 1B). Reverse-transcriptase polymerase chain reaction assays followed by Sanger sequencing determined that the *Lgr3^ag1^* deletion leads to readthrough from exon 7 directly into intron 7 and usage of an aberrant splicing donor within intron 7 directly into exon 9 (Fig. 1B and Supplementary Fig. 2A-D). The resulting transcript therefore has an intron-encoded premature termination codon (PTC) that truncates the Lgr3 protein one aminoacid after D_326_ (Fig. 1C). We conclude that *Lgr3^ag1^* encodes a severely compromised protein that is unlikely to bind ligand or signal due to a truncated ligand-binding LRR domain^5^, and absence of the seven transmembrane (7TM) domains and G protein coupling Carboxy terminus (Fig. 1C).

## Lgr3 couples growth to developmental timing

We noticed increased FA in *Lgr3^ag1^* adult wings. FA indexes (FAi) are increased by an order of ∼3 when compared to their *Lgr3^+/+^* controls [*p = 0.0028*, F-test for wing area FAi (Fig. 1D)]. This phenotype is indicative of uncoordinated imaginal disc growth during the larval stage, when wing size is determined, and is reminiscent of the increased FA phenotype of animals lacking the insulin/relaxin-like peptide Dilp83. We thus hypothesized that Lgr3 and Dilp8 act on the same pathway to promote developmental stability by coupling imaginal disc growth to developmental timing.

To test the hypothesis that Lgr3 and Dilp8 are in the same pathway, we asked the question whether *Lgr3* mutants show a similar defect as *dilp8* mutants in the ability to delay the onset of metamorphosis in response to induced abnormal tissue growth^3,4^. We did this by inducing tissue damage in an *Lgr3^ag1^* background and carried out pupariation timing assays. Tissue damage was induced by placing the proapoptotic gene *reaper* (*rpr*) under control of the wing-pouch *Beadex-Gal4* (*Bx>*)^3,9^, a combination that effectively caused a 72-*h* (median) delay in pupariation in control *Lgr3^+/+^* animals (Fig. 2A). Strikingly, this delay was reduced by 50 *h* in *Lgr3^ag1^* mutants [∼80%, *p <* 0.01, post-hoc test after Conover for Kruskal-Wallis test (hereafter, Conover post-hoc test); Fig. 2A]. The extent of this rescue is in line with previously published results using *dilp8* mutants^3,4^. This result demonstrates that Lgr3, like Dilp8, is critical to mediate the communication between abnormally growing imaginal discs and the prothoracic gland, strongly indicating that Lgr3 and Dilp8 act on the same pathway.

**Figure 2:**
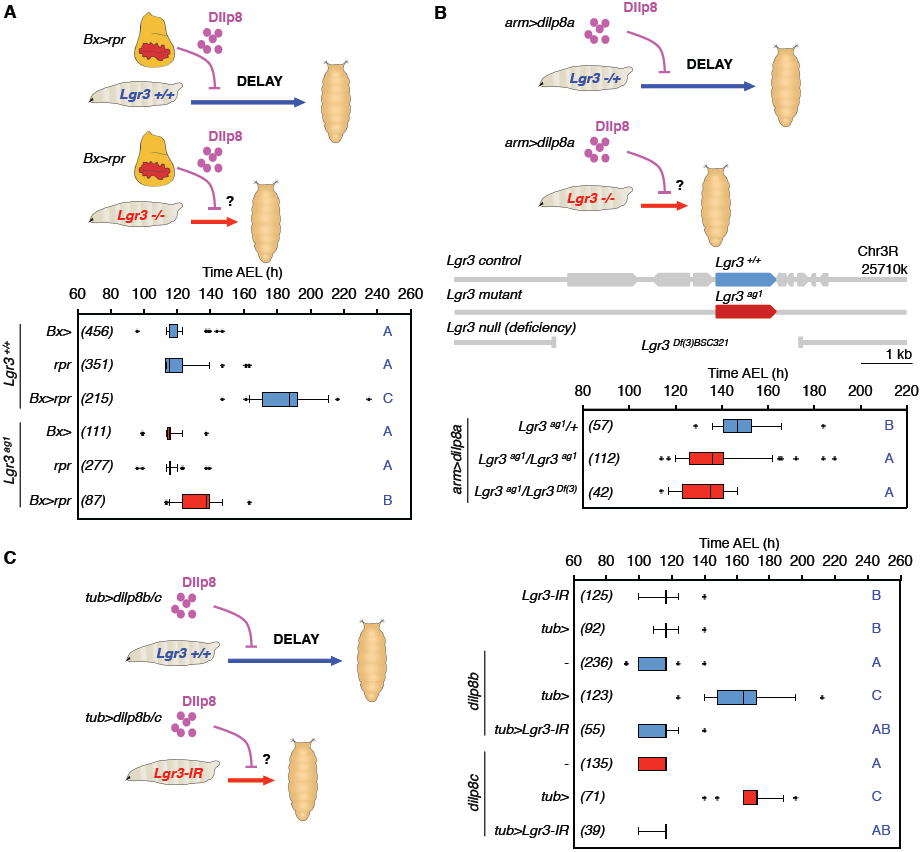
**Lgr3 couples imaginal disc growth to developmental timing by acting in the Dilp8 pathway. A,** Mutation of *Lgr3* abrogates interorgan communication between regenerating damaged discs (*Bx>rpr*) and the neuroendocrine centers coordinating the onset of metamorphosis. **B,** *Lgr3* acts in the Dilp8 pathway. Placing *Lgr3^ag1^* over a deficiency uncovering the *Lgr3* locus suppresses the delay caused by ectopic *dilp8* expression (*arm>dilp8a*). **C,** RNAi against *Lgr3* (*Lgr3-IR*) suppresses the delay caused by ectopic expression of either *dilp8b* or *dilp8c* transgenes^3^ under the control of the ubiquitous *tub>* driver. **A**, **B**, **C** Boxplots (see Methods) showing pupariation time [Time after egg laying (AEL) in *h*] of (*N*) larvae obtained from 6, 2, and 6 egg layings for panels **A**, **B** and **C,** respectively. Whiskers are 5 and 95% percentiles. Dots, outliers. *p <* 0.0001, Kruskal Wallis test for all panels. Genotypes sharing the same letter (blue) are not statistically different at alpha = 0.01, Conover post-hoc-test.

## Lgr3 acts in the Dilp8 pathway

To verify whether the Dilp8 activity is Lgr3 dependent, we tested if *Lgr3* was necessary for the developmental delay produced by ectopic Dilp8 expression in the absence of tissue growth abnormalities^3,4^. The delay in development induced by constitutive expression of a *UASP-dilp8::3xFLAG* (*UAS-dilp8a*) transgene under the control of the ubiquitous driver [using *armadillo-Gal4 (arm>)*] was suppressed in larvae homozygous for *Lgr3^ag1^* or trans-heterozygous for *Lgr3^ag1^* and a deficiency that completely uncovers the *Lgr3* locus (*Lgr3^Df(3)BSC321)^* (*p <* 0.01, Conover post-hoc test; Fig. 2B). These results show that animals lacking the *Drosophila* relaxin receptor Lgr3 are insensitive to ectopically produced Dilp8. Furthermore, as the suppression phenotype of *ag1* is indistinguishable from the *Df* allele, the results bring further evidence that *ag1* is a strong loss-of-function allele. To test whether *Lgr3* activity is required for the *dilp8-*dependent delay in a completely independent experimental setting, we induced a developmental delay using two previously-reported *dilp8* transgenes [*UAST-dilp8::3xFLAG* (*UAS-dilp8b and c*)]^3^ under the control of a ubiquitous driver, *tubulin-Gal4* (*tub>*), and reduced *Lgr3* activity by concomitant RNAi knockdown using a short hairpin [TRiP VALIUM22 (*Lgr3-IR-V22*)] line^10^, which reduces the steady-state *Lgr3* mRNA levels by ∼85% (*p=*0.043, Student’s t-test) (Supplementary Fig. 3A). Coherent with the mutant analyses, RNAi against *Lgr3* completely suppressed the *dilp8-*dependent delay (Fig. 2C). A second RNAi line producing a long hairpin against *Lgr3* [TRiP VALIUM10 (*Lgr3-IR-V10*)]^11^ also rescued the *dilp8-*dependent delay, albeit to a lesser extent than *Lgr3-IR-V22* (Supplementary Fig. 3B). This partial rescue was proportional to a weaker reduction in *Lgr3* mRNA levels (∼50%) than *Lgr3-IR-V22* (Supplementary Fig. 3A), suggesting that the Dilp8 delay is sensitive to *Lgr3* dosage. The *Lgr3-IR-V22* RNAi transgene was hereafter used in further experiments, as it phenocopies the *Lgr3* mutant phenotype best and gives the strongest RNAi reduction. Together with the *Lgr3* mutation analyses, the *Lgr3* RNAi experiments strongly place *Lgr3* in the Dilp8-dependent developmental delay pathway.

## Lgr3 is expressed in the CNS

To gain insight into the tissue and cellular expression pattern of *Lgr3* at the protein level, we used CRISPR/Cas9 mediated homologous repair^12,13^ to tag the endogenous Lgr3 protein at its Amino-terminus with superfolder green fluorescent protein (sfGFP*^14^*; Fig. 3A-B and Supplementary Fig. 4). We obtained one allele, *Lgr3^ag5^*, hereafter named *sfGFP::Lgr3*, which contained an intact *Lgr3* coding sequence downstream of the sfGFP insertion (Supplementary Fig. 4). sfGFP::Lgr3 labels ∼180 CNS cell bodies, consisting of ∼40 cell bodies in the brain proper and ∼140 in the ventral nerve cord (VNC) of 3^rd^ instar larvae CNS (Fig. 3C). No other larval tissue apart from the CNS had detectable fluorescence or anti-GFP staining. This rules out the possibility that *Lgr3* cell-autonomously controls ecdysone biosynthesis in the prothoracic gland downstream of Dilp8. The effectiveness of *sfGFP::Lgr3* as a protein translation reporter was further confirmed by analyzing alleles with premature termination codon-inducing (PTC+) indels in the *Lgr3* coding region following the inserted *sfGFP* sequence (Supplementary Fig. 4) or by using RNAi against endogenous *Lgr3* (Fig. 3C). Results show that either PTCs or RNAi against *Lgr3* effectively reduced sfGFP::Lgr3 expression to undetectable levels (Fig. 3C, right panels), demonstrating the reliability of the *sfGFP::Lgr3* endogenous protein reporter and further certifying the effectiveness of the *Lgr3* RNAi. Importantly, PTC+, but not PTC-alleles suppressed the delay induced either by raising the larvae in the presence of the genotoxic agent ethylmethanosulfonate (EMS), which induces apoptosis, tissue damage/regeneration, an imaginal-disc specific Dilp8 upregulation and consequentially a robust delay in the onset of metamorphosis^3^ (Fig. 3D) or by ectopically expressing Dilp8 (Supplementary Fig. 5). These results confirm that the PTC+ alleles are loss-of-function alleles and demonstrate that the PTC-*sfGFP::Lgr3 ag5* allele encodes a functional receptor. Together, these results strongly suggest that the Lgr3 protein acts in a subpopulation of CNS neurons.

**Figure 3:**
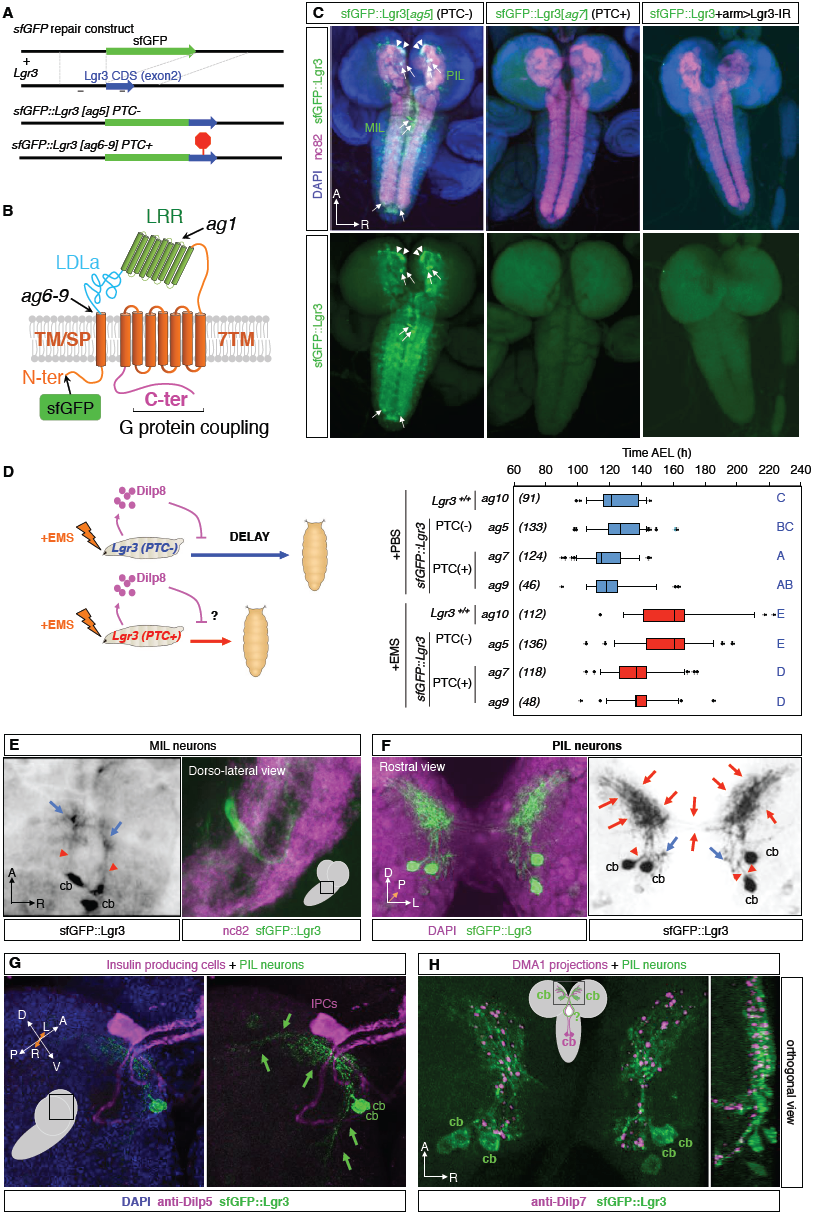
**Lgr3 is expressed in a subpopulation of CNS neurons. A**, CRISPR/Cas9-mediated sfGFP knock-in strategy used to generate the Lgr3 protein reporter allele *ag5* (named as *sfGFP::Lgr3*), which does not contain indels in the exon 2 region, and alleles *ag6-9* which contain PTC+ indels (Supplementary Fig. 4). Two thin black lines indicate the sites of the CRISPR gRNAs used. **B**, Lgr3 protein scheme depicting the approximate localization of the sfGFP insertion and the protein truncations caused by the PTC+ indel mutations. **C**, Sum of confocal z-stack slices of the CNS of a 3^rd^ instar larvae stained with anti-GFP (green) to show sfGFP::Lgr3 expression (green) and with anti-nc82 (magenta) and DAPI (blue) counterstains to show the synapses (neuropil) and nuclei, respectively. Arrows point to two bilateral pairs of PIL neurons (top), to the MIL neurons in the midline of the VNC (middle), and the distal VNC pair (bottom). Arrowheads point to the proximal projections of the PIL neurons. sfGFP::Lgr3 is also expressed in ∼170 other cell bodies, but at a lower level than in PIL and MIL neurons. No anti-GFP staining is detectable in the CNS of an animal carrying a sfGFP::Lgr3 insertion with a PTC+ indel [*ag7*] (middle panel) or in animals expressing RNAi against *Lgr3* (right panel). **D**, Boxplots (see Methods) showing pupariation time of (*N*) larvae obtained from 11 egg layings. Whiskers are 5 and 95% percentiles. Dots, outliers. *p <* 0.0001, Kruskal Wallis test. Genotypes sharing the same letter (blue) are not statistically different at alpha = 0.01, Conover post-hoc-test. **E**, MIL neuron cell bodies (cb) located deep in the VNC project ventrally and anteriorly (red arrowheads) for a short distance (red arrowheads). Adjacent anterior projections are depicted (blue arrows). **F**, Rostral view of the pars intercerebralis depicting the PIL neurons stained with anti-GFP [green (left) and black (right)] and counterstained with DAPI (magenta). Possible proximal dendritic arborizations are indicated with red arrows. Blue arrows indicate a ramification, likely axonic, with an undetermined terminus. Primary neurites (red arrowhead). **G**, PIL neurons associate closely to IPCs. Z-stack projection of confocal stacks stained with anti-GFP (green) and anti-Dilp5 (magenta). **H**, PIL neurons associate closely to Dilp7-producing DMA1 neurons in the pars intercerebralis. Z-stack projection and orthogonal view of confocal stacks stained with anti-GFP (green) and anti-Dilp7 (magenta).

Critically, we were also unable to detect sfGFP::Lgr3 expression in neurons directly innervating the ring gland (Fig. 3C), ruling out the possibility that Lgr3 neurons control developmental timing by direct cellular contact or synapsis with the ring gland, such as through PTTH-producing neurons^15^, the insulin producing cells (IPCs)^16^, or subesophageal serotonergic neurons^17^, all of which have been shown to modulate developmental timing. This scenario suggests that the Lgr3-positive neurons represent a novel distinct cellular population that has not been previously linked to growth and developmental timing control. To gain further insight into how Lgr3 neurons transmit the Dilp8 signal to the prothoracic gland, we looked into the neuroanatomy and neurotransmitter profile of *sfGFP::Lgr3* neurons in more detail. Notably, sfGFP::Lgr3 expression was not homogeneous in the ∼180 CNS cell bodies. It was most strongly expressed in a single pair of neurons in the very tip of the VNC, in a single dorso-ventral pair of midline neurons located deep in the thoracic segment of the VNC (hereafter abbreviated to MIL neurons, for midline internal Lgr3 neurons), and in a pair of bilateral neurons localized in the anterior part of the pars intercerebralis (hereafter abbreviated to PIL neurons, for pars intercerebralis Lgr3 neurons) (Fig. 3C, arrows and 3E) and (Fig. 3C, arrows and 3f). All three neuronal populations express *sfGFP::Lgr3* and present their characteristic neuroanatomy already at the L1 stage (Supplementary Fig. 6). Co-staining of *sfGFP::Lgr3* with *choline acetyltransferase (Cha)-Gal4* driving a *UAS-myr::tdTomato* reporter indicates that many *Lgr3* positive neurons are cholinergic, including all three major neuronal populations (PILs, MILs and the pair of distal VNC neurons (Supplementary Fig. 7). In contrast, co-staining with *glutamic acid decarboxylase (Gad1)-Gal4* and *Vesicular glutamate transporter promoter* (*VGlut)-Gal4*, which drive expression in GABAergic and glutamatergic neurons, respectively, label only very faintly the PIL neurons and the pair of distal VNC neurons and show no detectable staining in MIL neurons (Supplementary Fig. 8). While MIL neurons project their neurites ventrally and then anteriorly towards the brain, branching close to the base of the brain (Fig. 3E), the PIL neurons extend a single neurite centripetally into the neuropil that branches dorsally into highly arborized termini and ventrally into a slightly less arborized projection that proceeds towards the subesophageal region with a yet undetermined terminal branch (Fig. 3F). We noticed that the proximal arborizations of PIL neurons project to a region of the pars intercerebralis known to harbor the IPCs and anterior neurites from Dilp7-producing dorsal medial (DMA1)^18^ neurons projecting from the VNC. To verify this assumption, we co-stained *sfGFP::Lgr3* brains with anti-Dilp5 or anti-Dilp7 as markers for IPCs or DMA1 neuron projections, respectively. Our results show that PIL neuron proximal arborizations project towards the nearby IPC-bodies reaching them from immediately underneath, while the IPCs neurites extend through the PIL neuron arborization (Fig. 3G). PIL neurons also show an intimate association with the axonal projections of the Dilp7-producing DMA1 neurons (Fig. 3H). These results provide anatomical and possible functional context for an anterograde neuronal input for PIL neurons (considering that proximal branches are typically dendritic)^19^.

## Lgr3 acts in the CNS to delay development

The expression pattern of the sfGFP::Lgr3 reporter suggests that disrupting receptor function exclusively in the CNS should be sufficient to suppress the Dilp8-dependent delay in the onset of metamorphosis. To test this, we crossed the *Lgr3-IR-V22* RNAi line to the panneuronal CNS driver *elav-Gal4* (*elav>*) and performed EMS assays (Fig. 4A). Reducing *Lgr3* function with RNAi in the nervous system using *elav>* leads to complete suppression of the EMS-induced delay, as also observed with a ubiquitous knockdown [using *armadillo-Gal4 (arm>)*] (Fig. 4A). The neuronal requirement of *Lgr3* was further confirmed by knockdown of *Lgr3* in *elav* cells in a context of ectopic expression of *dilp8* (Supplementary Fig. 9A), and by rescuing neuronal expression of *Lgr3* in an *arm>dilp8 Lgr3-IR-V22* context by inhibiting *Lgr3* RNAi exclusively in the CNS by expressing the Gal4 inhibitor, Gal80, under the control of the *elav* enhancer (*elav-Gal80*; Supplementary Fig. 9B). As expected from our neuroanatomical and neurotransmitter biosynthesis expression pattern studies, knockdown of *Lgr3* in cholinergic neurons using *Cha-Gal4* also significantly suppressed the EMS-induced delay (Fig. 4B). In contrast, knockdown of *Lgr3* in the ring gland using *phantom (phm)-Gal4* line did not lead to a significant rescue of the EMS-induced delay (Fig. 4C). We conclude that *Lgr3* is required in the CNS, likely in one or more of the ∼180 Lgr3-positive cholinergic neurons described above, to convey the Dilp8-dependent developmental delay.

**Figure 4:**
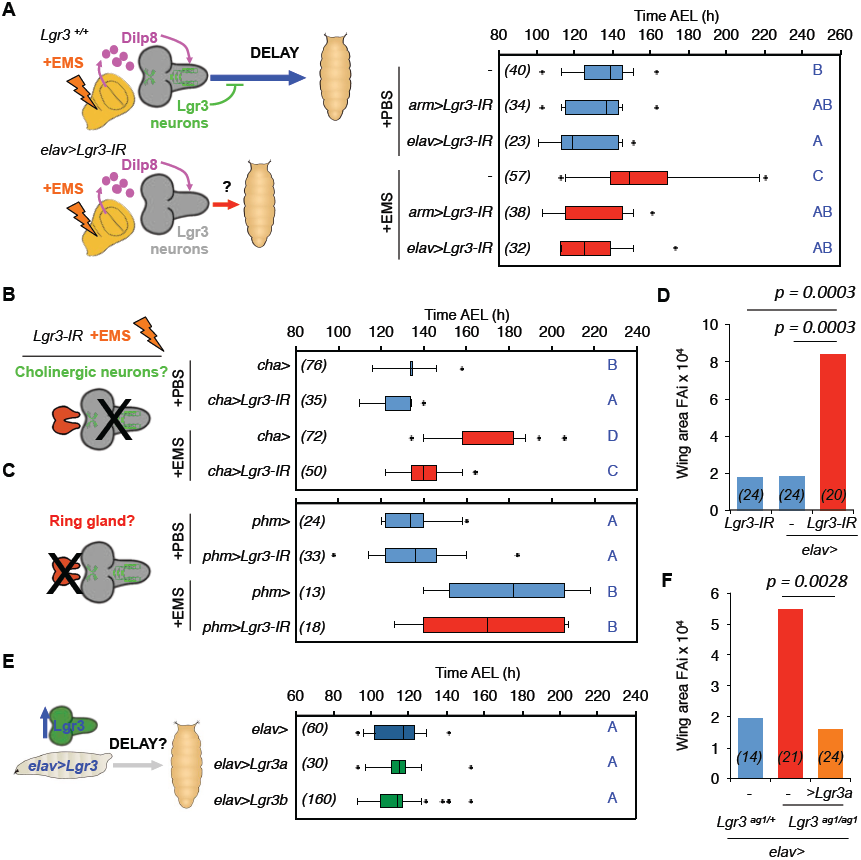
**Lgr3 is required in cholinergic neurons to couple growth and developmental timing. A,** CNS-specific RNAi of *Lgr3* rescues the EMS-induced delay. **B**, Removal of *Lgr3* in cholinergic neurons (using *cha>*) rescues the EMS-induced delay. **C**, Ring gland-specific RNAi of *Lgr3* does not significantly affect pupariation time in EMS assays. **D**, FAi in (*N*) animals expressing RNAi against *Lgr3* in neurons using *elav>* driver (F-test). **E**, Ectopic expression of *Lgr3* is not sufficient to delay the onset of metamorphosis. Two *UAS-Lgr3* transgene insertions (*a* and *b*) were tested. **F**, FAi in (*N*) animals carrying mutations in *Lgr3* and rescued with neuronal expression of *Lgr3a*. **A,B,C,E,** Boxplots (see Methods) showing pupariation time of (*N*) larvae obtained from 6, 2, 4, and 6 egg layings for panels **A, B, C,** and **E,** respectively. Whiskers are 5 and 95% percentiles. Dots, outliers. *p <* 0.0001, Kruskal Wallis test for all panels, except for panel **E,** where *p = 0.0413*. Genotypes sharing the same letter (blue) are not statistically different at alpha = 0.01, Conover post-hoc-test.

If, as described above, neuronal *Lgr3* is required to convey the Dilp8-dependent signal that promotes imaginal disc size stability by providing extra developmental time in the presence of disc growth problems, then the removal of *Lgr3* only in the CNS should also lead to an increased FAi, similarly to the increased FAi found in animals carrying mutations in *Lgr3* (Fig. 1D) or *dilp8*^[3]^. Consistent with this hypothesis, we find that FAi is increased by a factor of ∼4 in *elav>Lgr3-IR-RNAi* compared to controls (*p <* 0.001, in both F tests) (Fig. 4D). These results provide further evidence that neuronal *Lgr3* is critical in the pathway controlling developmental stability.

According to heterologous studies in human cell lines *in vitro*^7^, *Lgr3* is a constitutively active receptor. If this were true *in vivo* and if Dilp8 activity would inhibit Lgr3 activity, we would expect that by removing Lgr3 activity we would see a delay, which is not the case (see controls in Fig. 2A, Fig. 3D and Fig. 4A). To look at the effect of removing Lgr3 activity on pupariation timing at an increased resolution, we resynchronized larvae at the second to the third instar molt and confirmed that *Lgr3* mutants do not have a delayed development, but rather pupariate ∼4 h earlier than controls, independent of their genetic background (Supplementary Fig. 10). Even though our loss-of-function studies do not rule out constitutive activity, they are more consistent with a scenario where Lgr3 is activated either by Dilp8 or by other downstream Dilp8-dependent signals. We thus hypothesized increased Lgr3 levels would not lead to a delay. Alternatively, if Lgr3 were somehow constitutively active, it would delay pupariation timing if ectopically expressed. To conduct this experiment, we constructed a *UASP-Lgr3* transgene and expressed it in the CNS under the control of *elav>*. We find that the pupariation time profile of *elav>Lgr3* animals is indistinguishable from controls (Fig. 4E). Control experiments suggest that the *UASP-Lgr3* transgene carries functional Lgr3 activity, as it rescues the FAi of *ag1* animals when driven by *elav>* (Fig. 4F). These results argue against a simple explanation where Lgr3 is constitutively active^7^. Instead, they are in line with our findings that, in the absence of tissue growth aberrations or ectopic Dilp8 expression, Lgr3 activity does not have a major impact on timing the onset of metamorphosis. These results are coherent with a model where Lgr3 is activated either directly or indirectly by the Dilp8 signal and hint towards the existence of a dedicated tissue stress-sensing pathway in *Drosophila*.

## Lgr3 neurons relay the Dilp8 delay signal

To further narrow down the identity of the neurons requiring *Lgr3*, we tested the ability of two Janelia Gal4 lines^20^ carrying regulatory regions of the *Lgr3* locus, *GMR17G11-Gal4 (GMR17G11>)* and *GMR19B09-Gal4 (GMR19B09>)* (Fig. 5A), selected based on their larval brain expression patterns^21^, to suppress the EMS-induced developmental delay when driving *Lgr3-IR-V22.* While *GMR17G11>Lgr3-IR-V22* had no effect on the EMS-delay (Supplementary Fig. 11), the *GMR19B09>Lgr3-IR-V22* condition significantly reduced the 31.6-h EMS-dependent delay in pupariation time to 15.7 h (∼50% reduction, *p < 0.01*, Conover post-hoc test; Fig. 5B). We conclude that the subset of neurons expressing *GMR19B09>* is critical for the Dilp8 and Lgr3-dependent coupling of growth and developmental timing. Silencing of these same neurons by expression the inward rectifying K+ channel (Kir2.1) under the control of the *GMR19B09>* driver throughout embryonic and larval development was compatible with developmental progression and led to a significant suppression of the *EMS-*dependent delay (*p < 0.01*, Conover post-hoc test (Fig. 5C). These results strongly suggest that the activity of *GMR19B09>* neurons dictates the central Dilp8 and Lgr3-dependent response to tissue damage. To further characterize the neuroanatomy of *GMR19B09>* cells, we placed this driver together with a UAS-*myr::tdTomato* reporter transgene in a *sfGFP::Lgr3* background and carried out immunofluorescence assays of dissected CNSs. *GMR19B09>* drives expression in ∼270 neurons, ∼30 of which populate each brain hemisphere^21^. We find sfGFP::Lgr3 expression in ∼10 *GMR19B09>myr::tdTomato-*labeled neurons per hemisphere, two of which are the bright PIL neurons (Fig. 5D). We find no detectable *GMR19B09>myr::tdTomato* expression in MIL neurons and only a very faint trace, if any, in the posterior distal VNC pair (Supplementary Fig. 12). Of the ∼200 *GMR19B09-*positive VNC cells, roughly 60 co-express *sfGFP::Lgr3* at low, yet detectable levels. Any of the ∼70 CNS neurons coexpressing *sfGFP::Lgr3* and *GMR19B09>* are good candidate cells to convey the Dilp8 signal.

**Figure 5:**
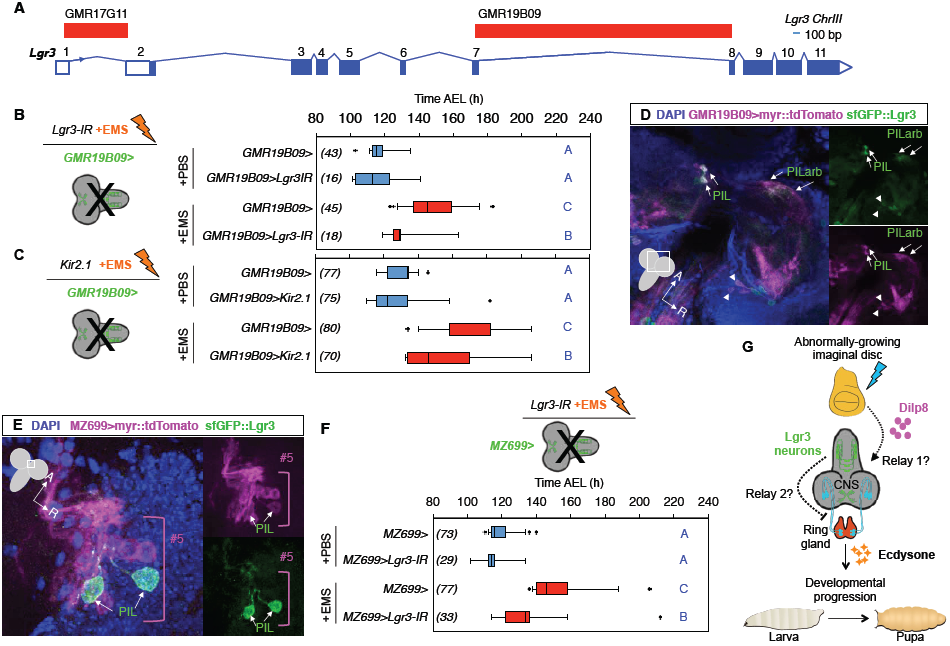
**A restricted subset of Lgr3 neurons relays the Dilp8 signal to the ring gland. A**, Scheme of the *Lgr3* locus depicting the regulatory elements^20,21^ tested in this study. **B**, RNAi of Lgr3 in *GMR19B09>* neurons rescues the EMS-induced delay. **C**, Silencing of *GMR19B09>* neurons by Kir2.1 expression suppresses the EMS-induced delay. **D**, *GMR19B09>* (magenta) drives expression in sfGFP::Lgr3-positive neurons (green): PIL neurons (PIL, arrows); PIL proximal arborizations (PILarb, arrows), and other neurons (arrowheads). **E,** PIL neurons (green) are a subset of #5 neurons (magenta) defined by *MZ699>*. **F**, RNAi of Lgr3 in *MZ699>* neurons rescues the EMS-induced delay. **G**, Cartoon depicting the Dilp8-Lgr3 abnormal tissue growth-sensing pathway. **B**,**C,F,** Boxplots (see Methods) showing pupariation time of (*N*) larvae obtained from 2 egg layings for each panel. Whiskers are 5 and 95% percentiles. Dots, outliers. *p <* 0.0001, Kruskal Wallis test for all panels. Genotypes sharing the same letter (blue) are not statistically different at alpha = 0.01, Conover post-hoc-test.

Since *GMR19B09* clearly colocalizes with PIL neurons, one of the cell populations that most strongly expresses *Lgr3*, we looked for other Gal4 lines that could allow the genetic manipulation of these neurons. We noticed that the *MZ-699-Gal4* (*MZ699>*) line drives expression in similar type of neurons, named #5 neurons^22^. Like PIL neurons, #5 neurons contribute to projections at the anteriormost region of the brain neuropil and to the median bundle tract that follows the esophageal foramen^22^. Immunofluorescence analyses of dissected CNS of larvae carrying *MZ699>myr::tdTomato* and *sfGFP::Lgr3* transgenes demonstrate that PIL neurons are a subset of #5 neurons (Fig. 5E). To test whether silencing *Lgr3* in *MZ699>* cells could rescue the tissue-damage-dependent developmental delay, we drove *Lgr3-IR-V22* under the control of *MZ699>* and exposed the larvae to EMS. We find that *MZ699>Lgr3-IR-V22* strongly suppresses the EMS-dependent delay (Fig. 5F). Apart from #5 neurons, colocalization of both transgenes in the brain region was only detected in 3 other neurons that appear to be a subset of #2 neurons, defined as medial antennocommissural tract relay interneurons whose fibers arborize in the laternal horn region^22^. However, *GMR19B09>* does not appear to label these cells, making it unlikely that they contribute to the Lgr3-dependent Dilp8 signal transduction. Other subesophageal neurons and many other VNC neurons, with the notable exception of the strong distal VNC neuronal pair, express both *MZ699>myr::tdTomato* and *sfGFP::Lgr3*. As *GMR19B09*> labeling of the subesophageal region is sparce^21^, and neither *MZ699>* or *GMR19B09>* drives detectable expression in MIL neurons (Supplementary Fig. 12 and Fig. 13), this suggests that either the PIL neuron expression overlap could be functionally relevant or that the Dilp8 signal requires Lgr3 expression in any of the ∼70 *sfGFP::Lgr3* positive VNC cells.

## Discussion

Different organs need to sense growth perturbations in distant tissues to coordinate their size and differentiation status during development^1^. Here, we have determined that the sensing of peripheral growth perturbations requires a novel population of CNS neurons expressing the Lgr3 relaxin receptor. Neuronal Lgr3 is required for the transmission of the peripheral growth aberration signal, Dilp8, to the prothoracic gland, which controls the onset of metamorphosis and thereby the cessation of imaginal disc growth^1,3,4,9,23^ (Fig. 5G). This work reveals a new Dilp8-Lgr3 pathway that is critical to ensure developmental stability in *Drosophila*. Our study opens many questions for further research, such as the determination of whether or not the interaction between Lgr3 and Dilp8 is direct, which of the ∼70 Lgr3-positive neurons are activated during Dilp8 expression, and how these connect to the ring gland. In this sense, the neuroanatomy of the sfGFP::Lgr3+ cells which also express *GMR19B09>* poses a problem of accessibility: if the activation of Lgr3 neurons by Dilp8 is direct, Dilp8 would have to somehow make its way across the blood-brain-barrier to activate these neurons deep in the brain. Alternatively, the Dilp8 signal could be received by other cells (either glial cells or other neurons with projections exposed to the hemolymph), and relayed through one or more steps before reaching the Lgr3+ cells (Fig. 5G, “Relay1?”). Furthermore, our study strongly suggests that the Lgr3+ cells that express both *MZ699>* and *GMR19B09>* are critical for the tissue damage developmental delay response. The fact that we were unable to detect *GMR19B09>myr::tdTomato* expression in the ring gland or in neurons innervating the ring gland strongly suggests that the *elav-*positive Lgr3+ neurons that are required for the Dilp8-dependent delay do not connect directly to the ring gland. Hence, it is likely that the Lgr3 neurons also need to relay the tissue stress signal at least once to the ring gland, either by secreting a second factor or connecting to a ring-gland innervating neuron (Fig. 5G, “Relay2?”). Together, these results indicate that the peripheral Dilp8 tissue damage signal is transduced through multiple steps before it reaches the ring gland, revealing unprecedented complexity and providing both important functional insight into the transduction of the Dilp8-dependent aberrant tissue growth signaling pathway, and opening fertile ground for further research.

The similarities between the neuroendocrine mechanisms controlling the larval to pupal transition in *Drosophila* and the hypothalamic-pituitary axis in vertebrates has been highlighted^2,24^. The neurosecretory cell-rich pars intercerebralis, in which the Lgr3-expressing PIL neurons are located, has anatomical, developmental and functional analogies to the hypothalamus, the structure that integrates the vertebrate CNS to the endocrine system via the pituitary gland. Similarly, the *Drosophila* pars intercerebralis connects the CNS to the endocrine ring gland complex via neurosecretory cells^25^, such as the IPCs^16,25^. Both systems have roles in stress-response, energy metabolism, growth, water retention, and reproduction^5,24^. The neuroanatomy of Lgr3-positive neurons, such as the PIL neurons (Fig. 3F-H), suggests they are well positioned to relay signals or to modulate the activity of ring gland-innervating neurons during tissue stress events that trigger Dilp8 secretion from the periphery. Candidate neurons that could interact with PIL neurons are the IPCs, PTTH neurons and DMA1 neurons, just to cite a few. Apart from arborizing in the pars intercerebralis region (Fig. 3F-H), PIL neurons send projections via the median bundle to the subesophageal region^21,22^. This region is known to harbor the serotonergic SE0-PG neurons, which directly innervate the PG, thereby regulating developmental timing as a response to nutritional cues^17^. It will be interesting to test whether PIL and SE0-PG neurons synapse in the subesophageal region and whether the latter also have a role in the tissue damage response.

As the timing of vertebrate developmental transitions, such as puberty, can also be altered by intrinsic and extrinsic factors affecting body growth, such as inflammatory disease and nutritional status^2^, the exploration of the role of relaxin signaling in modulating the hypothalamic-pituitary axis is a promising area for research. This is highlighted by the fact that the hypothalamus expresses relaxin receptors, including the Lgr3-homologue, RXFP1, in mammals and fish^5,26^, suggesting that a central neuroendocrine role for relaxin receptors might have evolved before the vertebrate and invertebrate split. A candidate peptide to regulate hypothalamic-pituitary stress-responses via relaxin receptors is the neuropeptide Relaxin-3 (RLN3), which has been traditionally viewed as being the ancestor ligand for all vertebrate relaxins^27,28^. RLN3 is strongly expressed in stress-responsive neurons from the nucleus incertus that directly innervate and modulate hypothalamic activity^5,29^-^31^. Our results therefore reveal an unexpected and striking similarity between the Dilp8-Lgr3 pathway and the vertebrate relaxin signaling pathway and hint to an ancient stress-responsive pathway coordinating animal growth and maturation timing.

## Acknowledgments

We thank C. Mirth, C. Ribeiro and A. Jacinto for their comments and suggestions; I. Miguel-Aliaga, A. Jacinto, P. Leopold, P. Domingos, C. Mirth, R. Teodoro, M. Dominguez, and M.L. Vasconcelos, for reagents. Stocks obtained from the Bloomington Drosophila Stock Center (NIH P40OD018537) were used in this study. AMG, FH, AM, AMD and TK are supported by the FCT, under the FCT Investigator Programme and FCT fellowships SFRH/BPD/94112/2013, PD/BD/52421/2013, SFRH/BD/94931/2013 and SFRH/BPD/74313/2010, respectively. AG is supported by the CONICET and UNS and YAV holds a CONICET fellowship. The work in the laboratory of AMG is funded by the CEDOC and the European Commission FP7 (PCIG13-GA-2013-618847). AG thanks NP Rotstein and LE Politi for providing funds and space to develop part of this project in their lab. The authors declare no competing financial interests.

## Author contributions

AG, FH, APC, AM, CN, MG, AMD, YV, TK and AMG performed experiments, analyzed data and contributed intellectually to the paper. AG, FH, APC, CN, TK and AMG were involved in pupariation time assays. FH, APC, AM, MG and AMD were involved in the generation and molecular characterization of mutants and transgenes. AG, APC, AM, YV and AMG were involved in FAi measurements. FH, APC, and AMG were involved in imaging. AG and AMG wrote the paper with the help of all authors.

## Methods

## Stocks

The *Drosophila melanogaster* stocks *w*^1118^ *Bx-gal4 (w^1118^ P{GawB}BxMS^1096^*), *UAS-rpr (w^1118^; P{UAS-rpr.C}14)*, *Lgr3-IR-V10 (y^1^ v^1^;{TRiP.JF03217}attP2)*, *Lgr3-IR-V22 (y^1^ sc* v^1^; {TRiP.GL01056}attP2/TM3 Sb^1^)*, *elav-Gal4 (P{w[+mW.hs]=GawB}elav^C155^), elav-Gal4 (P{w[+mC]=GAL4-elav.L}2/CyO), w^1118^;Mi{ET1}lgr3^MB06848^, w^1118^; sna^Sco^/SM6a, P{w[+mC]=hsILMiT}2.4, tub-Gal4 (y^1^ w*; P{tubP-GAL4}LL7/TM3, Sb[1])*, *w*^1118^; *Lgr3^Df(3)BSC321^/TM6C Sb^1^ cu^1^*, *Gad1-Gal4 (P{w[+mC]=Gad1-GAL4.3.098}2/CyO)*, *Cha-Gal4 (w^1118^; P{w[+mC]=Cha-GAL4.7.4}19B/CyO P{ry[+t7.2]=sevRas1.V12}FK1), GMR19B09-Gal4 (w^1118^; P{y[+t7.7] w[+mC]=GMR19B09-GAL4}attP2)*, and *GMR17G11-Gal4 (w[1118]; P{y[+t7.7] w[+mC]=GMR17G11-GAL4}attP2)* were obtained from the Bloomington Drosophila Stock Center at Indiana University. The stock *y*^1^ *M{w[+mC]=Act5C-Cas9.P}ZH-2A w\** (reference #*13)* was obtained from Bestgene. The stock *arm-gal4 (w*; P{w[+mW.hs]=GAL4-arm.S}11)* was a gift from P. Domingos. The stocks *UAS-dilp8b::3XFLAG and UAS-dilp8c::3XFLAG* (reference #3) were a gift from M. Dominguez. The stock *w*; P{GawB}Mz699* and *UAS-myr::tdTomato/CyO; TM2/TM6B* were a gift from M.L. Vasconcelos. Balanced lines were generated by crosses to the stock *w*^1118^; *If/CyO; MKRS/TM6B*, which was a gift from A. Jacinto. Stocks are maintained at low densities at 18°C in a 12-h light:dark cycle.

## Generation and molecular characterization of the *Lgr3^ag1^* allele

We generated a mutation in the Drosophila *Lgr3* locus by using the MiET1 transposase to remobilize the MB Minos element *Lgr3^MB06848^* (reference #*8*), which is inserted in the seventh *Lgr3* intron, <100 bp from exon 7 (Fig. 1B and Supplementary Fig. 1A). Virgin *Lgr3^MB06848^* females were crossed with males carrying the heat shock-inducible MiET1 transposase on a *Cy* balancer chromosome (*w*^1118^; *If/Cy-Minos; MKRS/TM6B*) and transferred to new vials every day^8^. After 48 h of development and until pupariation, the F1 progeny was given a 1-h 37°C heat-shock daily to induce the expression of the MiET1 transposase. *Cy-Minos/+; Lgr3^MB06848^/MKRS* or *TM6B* male adults were selected and individually crossed to the balancer strain *w*^1118^; *If/CyO; MKRS/TM6B*. A single eGFP-negative (lacking the *Mi{ET1} element) male was selected from each cross and mated with *w*^1118^; If/CyO; MKRS/TM6B* females. The putative *Lgr3* excisions were balanced over *TM6B* to obtain the following genotypes *w*^1118^; *+/+; Lgr3*^*MB06848*^ *excision/TM6B*. We obtained one imprecise excision that generated the *Lgr3^ag1^* mutant allele and three precise excisions named *Lgr3^ag2^, Lgr3 ^ag3^*, and *Lgr3^ag4^* all of which behaved the same way. In this study, the *Lgr3^ag2^* line was used as the genetic background control for the imprecise excision allele *Lgr3^ag1^*.

To molecularly characterize the *Lgr3^ag1^* deletion, we performed a series of polymerase chain reaction assays with primer pairs located around the *Lgr3^MB06848^* insertion (Supplementary Fig. 1A). *Lgr3^ag1^* was initially detected by the failure to amplify a 866-bp PCR product flanking the *Lgr3^MB06848^* insertion (green arrows, Supplementary Fig. 1A), and then a 240-bp PCR product using a pair of primers on exon 8 and 9 to the right of the *Lgr3^MB06848^* element (red arrows, Supplementary Fig. 1A, B), indicating that a deletion occurred downstream of the element following its remobilization. A PCR using a primer pair in the exon 11 produced a positive result, indicating that exon 11 was present in *Lgr3^ag1^* (not shown). This suggested that while the *Lgr3^ag1^* deletion was very large, its breakpoints were confined within the *Lgr3* locus. We then tried to amplify a 5.3-kb PCR product with a primer upstream of the *Lgr3^MB06848^* position (green forward arrow, Supplementary Fig. 1A-C) and another on exon 11 (purple arrow, Supplementary Fig. 1A-C). Instead of the predicted >5.3 kb product, we obtained a ∼1.6 kb product (Supplementary Fig. 1C), demonstrating that *Lgr3^ag1^* is a large deletion of approximately 3.8 kb in the *Lgr3* locus (Supplementary Fig. 1D). This alone indicates that at least 69 aminoacids (aa; I_327_-T_395_) are lacking in the *Lgr3^ag1^* Leucine Rich Repeat (LRR) domains, which are critical for relaxin ligand binding in vertebrate relaxin receptors^5^.

Next, we performed reverse-transcriptase (RT)-PCR analyses with mRNA isolated from *Lgr3^ag1^* and controls *Lgr3^+/+^* and the original *Lgr3^MB06848^* stock. *Lgr3^ag1^* generated a smear with a major band that was ∼200 bp smaller than the other control genotypes (Supplementary Fig. 2A; Supplementary Fig. 2B, C are RT- and *rp49* control reactions). Sanger sequencing determined that the *Lgr3^ag1^* deletion leads to readthrough from exon 7 directly into intron 7 and usage of an aberrant splicing donor within intron 7 directly into exon 9 (Supplementary Fig. 2D). The resulting transcript therefore has an intron-encoded premature termination codon (PTC) that truncates the Lgr3 protein one amino-acid after D_326_ (Fig. 1E). We conclude that *Lgr3^ag1^* encodes a severely truncated protein without the seven transmembrane (7TM) domains and G protein coupling Carboxy terminus (Fig. 1E).

## Generation of *pUASP-Lgr3* and *pUASP-dilp8* flies

A *Drosophila* and *Homo sapiens* codon-optimized cDNA corresponding to full-length *Lgr3* was synthetized *de novo* and cloned into *pUASP*^32^ using KpnI and NotI sites to make *pUASP-Lgr3.* This plasmid was injected into *w*^*1118*^ and two independent insertions were tested, *pUASP-Lgr3a (M3, on Chr III)* and *pUASP-Lgr3b (M6, on Chr II).* A similar protocol was used to place the *dilp8::3xFLAG* construct described in reference #3 into *pUASP*, making *pUASP-dilp8a.*

## *Generation of sfGFP::Lgr3* and PTC-inducing alleles

We used a CRISPR/Cas9 mediated homologous repair strategy^12^,^13^ to tag the endogenous Lgr3 protein at its Amino (N)-terminus with superfolder green fluorescent protein (sfGFP)^14^ followed by a flexible linker spacer sequence GSGSGS (Fig. 3A and Supplementary Fig. 5). The following guide RNAs (gRNA1 and gRNA2, designed with http://tools.flycrispr.molbio.wisc.edu/targetFinder/)^12^ were synthetized and cloned into pU6-BbsI-chiRNA:

gRNA 1

Fw CTTCGGAGCACTCAATTCCCACTC (CGG)

Rv AAACGAGTGGGAATTGAGTGCTCC

gRNA 2

Fw CTTC<u>G</u>CAAACTCAAGTAGAATATCA (CGG)

Rv AAACTGATATTCTACTTGAGTTTG<u>C</u>

The PAM regions (blue) are located right after the forward primers of both gRNAs 1 and 2, as shown above. Red nucleotides are used for cloning purposes.

As a repair cassette we designed the following sequence, where each color represents the following:

**Yellow: Homology region up to *Lgr3* ATG site**

**Red: *Lgr3* ATG**

**Green: sfGFP**

**Blue: Spacer (GSGSGS)**

**Violet: Homology region after *Lgr3* ATG site**

CACTTAAAACTCTTCTCCGCGAGCTGTGAACATTAGCCAAATGAAGTGACAAGAAATTAACGCAAAAATAAAACAAGAAGACGGAGCGGTATAAGAAATAATAATATAAAAACTCAATGAGTCAGCACCGCATCAGCTCCTGCTGCTGTTGTTCTTCTTATTGCTGTTGTTTGTGGGGGCGTGGCCGGAGTGGGAATTGAGTGCTCCTAATGATGAACTCGGTCAAGGAGCCAGTGCAGCCATGGTGGCCAAGTAATTAGATAAGCGAGCGTGCAAAACAGGAGCAAACCGATAAATCGCC<u>ATG</u>CGTAAGGGCGAGGAGTTGTTCACGGGAGTTGTGCCCATATTGGTTGAGCTGGATGGAGATGTGAATGGCCACAAGTTCAGTGTGCGGGGTGAGGGAGAAGGAGACGCAACAAACGGTAAGCTGACACTGAAGTTCATTTGTACTACGGGCAAGCTCCCGGTGCCATGGCCCACATTGGTCACCACCCTGACCTATGGCGTGCAATGCTTCGCCCGATATCCAGATCATATGAAGCAGCATGATTTCTTTAAGTCGGCCATGCCCGAGGGTTACGTACAAGAGCGCACTATTAGCTTTAAGGACGACGGTACGTATAAAACCAGGGCTGAGGTGAAGTTTGAGGGTGATACCCTGGTGAACCGCATTGAATTGAAGGGCATCGATTTTAAGGAGGACGGCAACATCCTGGGCCACAAGCTCGAATATAATTTTAATAGCCATAATGTTTACATTACCGCGGACAAGCAGAAGAATGGAATTAAGGCTAATTTCAAGATCCGACATAATGTGGAGGACGGATCCGTTCAGTTGGCCGATCACTACCAGCAAAACACCCCCATCGGAGATGGCCCCGTCCTGCTGCCCGATAACCACTACCTGAGTACCCAGTCCGTCCTGTCGAAGGATCCTAATGAGAAGCGGGATCATATGGTGCTGCTGGAGTTTGTGACTGCCGCCGGCATAACGCATGGAATGGACGAGCTGTATAAA<u>GGCTCCGGTAGTGGTTCC</u>GTCTACGGCAGGAGCATCGCCGTAGGCTTCTGTCTGATGACCGTCGTCCTTCTGCTGGCCGCCGTGATATTCTACTTGAGTTTGGGTGAGTCCTTAGAGTGATGTCCTTTCAAAATTCCATCATTCGCAAACCTAAATAATTTCTGAATCAAGAATGTTCAAAATCTTAGCAATTATTATACGCATAATTTGTGAAACTACTTAAAGTTCTTTTAAAACTTGAGCTGCTGTAAATTTCTATATATACTTTCGTATCCTTAAAGGGTTCCTTCGCTTGAAGCAAAAACCAAAATCAAATTCCAAACTGCAAA

The repair cassette was synthetized *de novo* into a pUC57 plasmid and co-injected with the two gRNAs into the stock *y[1] M{w[+mC]=Act5C-Cas9.P}ZH-2A w[*]*, which strongly and ubiquitously expresses a human codon optimised cas9 (reference #*12)*. The adults originating from this injection were separately crossed to a *MKRS/TM6B* stock. Males from each F1 cross were separately crossed again to MKRS/TM6B. When larvae appeared in the vials, we extracted gDNA from the F1 male progenitor and made PCR reactions using the primer pairs:

Lgr3_crispr_testF - CCAATAACTTTAAGCCGTCTGTG

SuperfolderGFP_R - CAGCACCATATGATCCCGCT

SuperfolderGFP_F - CCTATGGCGTGCAATGCTTC

Lgr3_crispr_testR - TATAGCTGTGCGAATTTCTCGAT

These primers were designed to indicate the correct insertion of the sfGFP repair cassette from both sides in the genome. Positive hits were sequenced (*ag5-ag9*). Ten negative hits were also sequenced and two were retained as background controls (*ag10* and *ag11*). The *y*^*1*^ *M{w[+mC]=Act5C-Cas9.P}ZH-2A w\** cassette was removed by selecting against eye color and stocks were kept as homozygous stocks, except for allele *ag8*, which was kept balanced over TM6B.

## Measurement of the developmental timing of pupariation

Male and virgin female flies aged 3-10 days old were crossed and 1-2 days later transferred to an agar plate with yeast-sucrose paste (1:1). The next morning, the flies were transferred to a fresh plate to lay eggs for 3-6 h, depending on the experiment. To control for overcrowding, larvae were transferred to vials containing normal Drosophila food in 3-12 batches typically between 10-30 depending on the experiment. Survey of pupae consists in counting the number of pupae in each time interval. The final total n of pupae for each genotype in each experiment is depicted in the respective figures. For pupariation time assays, we aimed at obtaining at least 30 individual larvae per genotype, yet some genotypes were sick or difficult to obtain due to balancer chromosomes, yielding less larvae of the correct genotype to score. Male and females were scored together. The time of pupariation of each larvae was determined and genotypes were compared with the Kruskal-Wallis non-parametric test followed by post-hoc ranks test after Conover, with α = 0.01 or 0.05, as indicated in each figure, using the software Infostat. Data was plotted in box-plots representing median, 25 and 75% quartiles, and 5-95% percentiles as whiskers. Data points falling outside of the 5-95% interval were plotted as outliers. The Kruskal-Wallis test is a rank-based nonparametric test that can be used to determine statistically significant differences between two or more groups. It is a nonparametric alternative to the one-way ANOVA, and as such, normal distribution is not required. Developmental timing assays were performed at 25°C under constant light, except for experiments reported in Fig. 2A, and Supplementary Figure 3B, which were done in a 12-h light:dark cycle, and EMS experiments (see below), which were done in the dark to minimize the reaction of EMS with light.

## Duration of the third instar

Egg collections were performed on normal food plates and larvae were reared at controlled densities without additional yeast (about 200 eggs/60 mm diameter normal fly medium plate). Newly molted third instar larvae were collected every 2 h as described previously^33^, ^34^. After staging, collected larvae were raised in a normal cornmeal/molasses medium at 20-30 larvae/vial without additional yeast. Pupariation time was observed every 2 h until all treated larvae pupariated or died. Male and females were scored together. We defined pupariation as cessation of movement with evaginated spiracles. These experiments were performed at 25°C under constant light.

## Ethyl methanesulfonate (EMS) treatment

Larvae were collected as described above and transferred 72 h after egg laying to fresh food with 10 mM of EMS (Sigma) or PBS as control. Developmental time was measured as indicated above. EMS food was prepared as follows: food was melted and cooled to 55°C, an appropriate volume of freshly made EMS stock solution in PBS was added and thoroughly mixed and 3 ml per tube were dispensed. EMS and PBS tubes were kept in the dark as much as possible throughout the experiments.

## Immunofluorescence staining

Brains of wandering 3^rd^ instar larvae or 1^st^ instar larvae were dissected in cold PBS, fixed for 30 min in 4% paraformaldehyde, rinsed with PBS with Triton (0.3%) (PBST), incubated with primary antibody overnight and with fluorescently labelled secondary antibody for 2 h. Nuclei were counterstained with DAPI (Sigma) and tissues mounted in Fluoromount-G (Southern Biotech). Antibodies used were: mouse anti-GFP 1:200 (DSHB, 12E6), mouse anti-nc82 1:250 (DSHB), rabbit anti-GFP 1:200 (Life technologies, A11122), rabbit anti-Dilp7 1:5000 (I. Miguel-Aliaga^18^), rat anti-Dilp5 1:400 (gift from P. Leopold^35^). Images were obtained with a Zeiss LSM 710 Confocal Microscope and images were analyzed using FIJI software^36^. Typically 5-10 brains were mounted for observation and one representative image per genotype is depicted in figures. Brains from male and female larvae were scored together.

## Adult wing measurements

Pairs of the left and right wings of male individuals reared at 29°C were rinsed with ethanol and mounted in a glycerol: ethanol solution. Photos were obtained in a Zeiss Axiovert 40 CFL microscope. The wing areas and wing lengths were calculated as previously described^3^, ^39^, using Fiji. We used the Fluctuating Asymmetry index (FAi) to assess intra individual size variations between the left and right wings^39^. Namely, FAi = Var(Ai), where Ai are the normalized differences between left and right wing areas of each individual Ai=A_left_-A_right_/[(A_left_+A_right_)/2]. Results were compared statistically using the F-test for the significance of the difference between the FAi of the samples, an appropriate test for dispersion^3^, ^39^. Bonferroni corrections (α/n) for multiple comparisons was applied to Fig. 4D and Fig. 4F, using α = 0.05

To control for measurement errors, we measured the area of the same wing three times. Values obtained were 18310.2 ± 30.5 AU, which gives a coefficient of variation (CV) of 0.17%, which is smaller than the CV of wing areas of control *Lgr3^+/+^* by a factor of ∼23 (CV = 3.88%, n = 48, flies reared at 29°C).

## gDNA, cDNA, RT-PCR and qRT-PCR analyses

gDNA was extracted from a group of flies or single flies^38^. Briefly, the flies were macerated using pellet pestles and homogenized in 100 μl DNA Extraction Buffer (1 M Tris-HCl at pH 8.2, 0.5 M EDTA, 5 M NaCl). Then, we added 1 μl Protease K 50 ng/μl (Roche), and incubated the mixture at 37°C for 1 h, followed by 95°C for 5 min, to inactivate the protease.

RNA was extracted using the Direct-zol(tm) RNA MiniPrep kit (Zymo Research), following manufacturer’s instructions. The material used for the RT-PCR experiments described in Supplementary Fig. 2 was 15 virgin males aged between 3-7 days were macerated using pellet pestles and homogenized in 500 μl TRI Reagent® and centrifuged at 12000 g for 1 min, to lower tissue debris. An extra DNAse treatment (Turbo DNA-free kit, Ambion, Life Technologies) was performed to reduce gDNA contamination. cDNA synthesis was performed using the Maxima First Strand cDNA Synthesis Kit for RT-qPCR (ThermoScientific), following manufacturer’s instructions. For this study, PCR and RT-PCR primers were designed and their specificity tested using Primer - BLAST or primer3. A T100 Thermal Cycler(tm) (BioRad) for performing the PCR steps. The following primers were used for PCR analyses described in Supplementary Fig. 1-2.

Lgr3_salto_fw (expected product size 866 bp)

Fw CCGACGCCTTGCTGCTAACT

Rv TTTATGGAGCGGGCGTGGTC

Lgr3_exonshort Lgr3 (expected product size 331 bp)

Fw CCGACGCCTTGCTGCTAACT

Rv GTGCGTTATGAGGTTGTGCTG

Lgr3_exon3p Lgr3 (expected product size 240 bp)

Fw CGCCTTGTCGGTAATCCCAT

Rv GTGGCTCCATTAAACTGCTGC

Lgr3_exons (expected product size 5307)

Fw CCGACGCCTTGCTGCTAACT

Rv CAAAGACCACCAACCAGGCGTA

*rp49* (control)

Fw TTGAGAACGCAGGCGACCGT

Rv CGTCTCCTCCAAGAAGCGCAAG

qRT-PCR experiments were performed using Lightcycler® 96 (Roche) using the FastStart Essencial DNA Green Master dye and polymerase (Roche). The final volume for each reaction was 10 μl, consisting of 5μl of dye and polymerase (master mix), 2 μl of cDNA sample, and 3 μl of the specific primer pairs. The following primers were used:

*rp49* (control)

Fw TTGAGAACGCAGGCGACCGT

Rv CGTCTCCTCCAAGAAGCGCAAG

*Lgr3*

Fw GCTGGGTGCCCATCATCGTTAT

Rv CAAAGACCACCAACCAGGCGTA

Primer efficiency for qRT-PCR was tested by serial dilution. Data was expressed as %*rp49* according to the formula: %*rp49* = (2^-(ΔCqLgr3-ΔCqrp49))*100. The geometric mean ± the standard error of the mean of three biological repeats was used and data was analyzed by one-tailed unpaired Student’s t-test using Bonferroni corrections (α/n) for multiple comparisons, using α = 0.05. This test assumes the independent samples have equal variances.

## General study design and statistics

In all experiments reported in this work, no data point was excluded. All data points, including outliers, are represented in the Figures and were used in the statistical analyses. No blinding was done and no particular randomization method was used to individuals to experimental groups.

## References

1. Shingleton, A. W. The regulation of organ size in Drosophila: physiology, plasticity, patterning and physical force. Organogenesis 6, 76–87 (2010).

2. Hariharan, I. K. How growth abnormalities delay **“**puberty **„** in Drosophila. Sci Signal 5, pe27 (2012).

3. Garelli A., Gontijo, A. M., Miguela, V., Caparros, E., Dominguez, M. Imaginal discs secrete insulin-like peptide 8 to mediate plasticity of growth and maturation. Science 336, 579–582 (2012).

4. Colombani, J., Andersen, D. S., Léopold, P. Secreted peptide Dilp8 coordinates Drosophila tissue growth with developmental timing. Science 336, 582–585 (2012).

5. Bathgate, R. A., et al., Relaxin family peptides and their receptors. Physiol Rev 93, 405–480 (2013).

6. Van Hiel, M. B., Vandersmissen, H. P., Van Loy, T., Vanden Broeck, J. An evolutionary comparison of leucine-rich repeat containing G protein-coupled receptors reveals a novel LGR subtype. Peptides. 34, 193–200 (2012).

7. Van Hiel, M. B., Vandersmissen, H. P., Proost, P. Vanden Broeck, J. Cloning, constitutive activity and expression profiling of two receptors related to relaxin receptors in Drosophila melanogaster. Peptides. 68, 83–90 (2015).

8. Metaxakis, A., Oehler, S., Klinakis, A., Savakis, C. Minos as a genetic and genomic tool in Drosophila melanogaster. Genetics 171, 571–581 (2005).

9. Halme, A., Cheng, M., Hariharan I. K. Retinoids regulate a developmental checkpoint for tissue regeneration in Drosophila. Curr Biol 20, 458–463 (Mar 9, 2010).

10. Ni, J. Q., et al. A genome-scale shRNA resource for transgenic RNAi in Drosophila. Nat Methods, 8, 405–407 (2011).

11. Ni, J. Q., et al. A Drosophila resource of transgenic RNAi lines for neurogenetics. Genetics, 182, 1089–1100 (2009).

12. Gratz, S. J., et al. Genome engineering of Drosophila with the CRISPR RNA-guided Cas9 nuclease. Genetics 194, 1029–1035 (2013).

13. Port, F., Chen, H. M., Lee, T., Bullock. S. L. Optimized CRISPR/Cas tools for efficient germline and somatic genome engineering in Drosophila. Proc Natl Acad Sci U S A, 111, E2967–E2976 (2014).

14. Pédelacq, J. D., et al. Engineering and characterization of a superfolder green fluorescent protein. Nat Biotechnol 24, 79–88 (2006).

15. McBrayer, Z. et al. Prothoracicotropic hormone regulates developmental timing and body size in Drosophila. Dev Cell 13, 857–871 (2007).

16. Rulifson, E. J., Kim, S. K., Nusse, R. Ablation of insulin-producing neurons in flies: growth and diabetic phenotypes. Science 296, 1118–1120 (2002).

17. Shimada-Niwa, Y., Niwa, R. Serotonergic neurons respond to nutrients and regulate the timing of steroid hormone biosynthesis in Drosophila. Nat Commun 5, 5778 (2014).

18. Miguel-Aliaga, I., Thor, S., Gould A. P. Postmitotic specification of Drosophila insulinergic neurons from pioneer neurons. PLoS Biol 6, e58 (2008).

19. Hartenstein, V., Spindler, S., Pereanu, W., Fung, S. The development of the Drosophila larval brain. Adv Exp Med Biol 628, 1–31 (2008).

20. Jenett, A., et al., A GAL4-Driver Line Resource for Drosophila Neurobiology. Cell Rep 2, 991–1001 (2012).

21. Li, H. H., et al., A GAL4 driver resource for developmental and behavioral studies on the larval CNS of Drosophila. Cell Rep 8, 897–908 (2014).

22. Ito, K., Sass, H., Urban, J., Hofbauer, A., Schneuwly, S. GAL4-responsive UAS-tau as a tool for studying the anatomy and development of the Drosophila central nervous system. Cell Tissue Res 290, 1–10 (1997).

23. Gilbert, L. I., Rybczynski, R., Warren, J. T. Control and biochemical nature of the ecdysteroidogenic pathway. Annu Rev Entomol 47, 883–916 (2002).

24. de Velasco, B., et al. Specification and development of the pars intercerebralis and pars lateralis, neuroendocrine command centers in the Drosophila brain. Dev Biol 302, 309–323 (2007).

25. Siegmund, T., Korge, G. Innervation of the ring gland of Drosophila melanogaster. J Comp Neurol 431, 481–491 (2001).

26. Donizetti, A., et al. Characterization and developmental expression pattern of the relaxin receptor rxfp1 gene in zebrafish. Dev Growth Differ 52, 799–806 (2010).

27. T. N. Wilkinson, R. A, Bathgate. The evolution of the relaxin peptide family and their receptors.Adv Exp Med Biol 612, 1–13 (2007).

28. Yegorov, S., Good, S. Using paleogenomics to study the evolution of gene families: origin and duplication history of the relaxin family hormones and their receptors. PLoS One 7, e32923 (2012).

29. McGowan, B. M., et al. Relaxin-3 stimulates the hypothalamic-pituitary-gonadal axis.Am J Physiol Endocrinol Metab 295, E278–E286 (2008).

30. Watanabe, Y., Miyamoto, Y., Matsuda, T., Tanaka, M. Relaxin-3/INSL7 regulates the stress-response system in the rat hypothalamus. J Mol Neurosci 43, 169–174 (2011).

31. McGowan, B. M., et al. Relaxin-3 stimulates the neuro-endocrine stress axis via corticotrophin-releasing hormone. J Endocrinol 221, 337–346 (2014).

32. Rørth, P. Gal4 in the Drosophila female germline. Mech Dev 78, 113–118 (1998).

33. Mirth, C., Truman, J. W., Riddiford, L. M. The role of the prothoracic gland in determining critical weight for metamorphosis in Drosophila melanogaster. Curr Biol 15, 1796–1807 (2005).

34. Koyama, T. Rodrigues, M. A., Athanasiadis, A., Shingleton, A. W., Mirth, C. K. Nutritional control of body size through FoxO-Ultraspiracle mediated ecdysone biosynthesis. Elife 3, (2014) doi: 10.7554/eLife.03091.

35. Geminard, C., Rulifson, E. J., Leopold, P. Remote control of insulin secretion by fat cells in Drosophila. Cell Metab 10, 199–207 (2009).

36. Schindelin, J., et al. Fiji: an open-source platform for biological-image analysis.Nat Methods 9, 676–682 (2012).

37. Mirth, C., Truman, J. W., Riddiford, L. M. The ecdysone receptor controls the post-critical weight switch to nutrition-independent differentiation in Drosophila wing imaginal discs. Development 136, 2345–2353 (2009).

38. Carvalho, G. B., Ja, W. W., Benzer, S. Non-lethal PCR genotyping of single Drosophila.Biotechniques 46, 312–314 (2009).

39. Palmer, A. R., Strobeck, C. Fluctuating Asymmetry: Measurement, Analysis, Patterns Annu Rev Ecol Syst 17, 391–421 (1986).

